# Video-based eye movements are linked to cortical and pupil-based arousal in human sleep

**DOI:** 10.64898/2026.06.10.731319

**Authors:** Manuel Carro-Domínguez, Stella Oberlin, Timona Leandra Oesch, Nicole Wenderoth, Sarah Nadine Meissner, Caroline Lustenberger

## Abstract

**Study Objectives:** To quantify high-resolution video-based eye kinematics across sleep macro- and microstructure and determine their coupling with pupil-based and cortical arousal markers.

**Methods:** We recorded polysomnography and utilized infrared video-based eye-tracking in 17 healthy adults. Computer vision was employed to extract eye position and speed, which were related to pupil size (subcortical arousal marker) and EEG spectral slope (cortical arousal marker) across sleep stages, rapid eye movement (REM) substates, and K-complexes.

**Results:** Eye kinematics varied significantly across sleep stages (p<0.001), except for horizontal pupil position (p=0.192). Video-based eye position and speed are sufficient to classify stages with above chance level accuracy (47%). Eye movement speed positively correlated with pupil size during non-REM sleep (R≥0.21, p<0.001) and with spectral slope during wakefulness and REM sleep (R≥0.11, p<0.018). These strengths of the correlations differ depending on the direction of the eye movement. Phasic REM exhibited faster eye movements, larger pupil size (p<0.001), and a flatter spectral slope (p=0.014) compared to tonic REM, indicating elevated subcortical and cortical arousal. K-complexes were accompanied by increased eye movement speed (p<0.05) and a steeper spectral slope (p≤0.010), suggesting a transient shift toward a sleep-protective cortical state despite concurrent oculomotor activation.

**Conclusions:** Video-based eye-tracking reveals that eye movements are quantitatively coupled to brain-wide arousal fluctuations in a state-dependent manner. This methodology provides a ground-truth framework for resolving fine-grained eye dynamics during sleep, offering a high-fidelity tool for sleep phenotyping and clinical assessment.

**Statement of significance:** Traditional sleep monitoring relies on low-resolution electrical signals that often confound true eye movements with brain or muscle activity. This study uses high-resolution video tracking to establish a physical ground truth for eye kinematics across human sleep. We demonstrate that eye movements are coupled with both cortical and subcortical arousal levels in a state-dependent manner, providing a clearer neurophysiological distinction between, for example, sleep substates like tonic and phasic REM. This framework reveals how the eyes serve as a non-invasive window into the brain’s internal arousal state. Such insights address critical gaps in understanding sleep microarchitecture and offer a novel pathway for developing high-fidelity biomarkers for neurological disorders, such as Parkinson’s disease, characterized by disrupted sleep-related arousal.

## Introduction

Arousal level fluctuations during sleep are a fundamental brain process reflecting a continuum of activation across cortical and subcortical networks that drive restorative and regenerative processes ^1–4^. These fluctuations can be indexed by complementary physiological proxies: the aperiodic component of the EEG power spectrum (spectral slope) as a marker of cortical arousal, and pupil size as a marker of pupil-based arousal linked to subcortical neuromodulatory systems^5–8^. Using these proxies, recent work has revealed distinct cortical and pupil-based arousal level dynamics across sleep macrostructure and within microstructural events^9^. As average cortical and pupil-based arousal levels decrease with deeper stages of sleep, pupil-based arousal levels fluctuate at an infraslow timescale within NREM sleep states^9^.

However, it remains unclear how these relatively novel arousal proxies relate to the traditional physiological markers that are foundational for sleep medicine to detect changes in brain state, specifically electrooculography (EOG)-derived eye movements. While EOG activity is known to decrease with deeper stages of sleep until rapid eye movement (REM) sleep is reached, the specific link between oculomotor dynamics and the underlying arousal state is complex. For instance, recent work utilizing the EEG spectral slope has further characterized REM sleep, subdividing it into ‘tonic’ periods of relative ocular quiescence and lower cortical arousal, and ‘phasic’ periods defined by bursts of saccadic eye movements and higher cortical arousal^10^.

This raises a critical question: are eye movements during sleep macro- and microstates merely descriptive labels, or do they result from specific changes in brain states driven by arousal level fluctuations? Animal research suggests the latter, demonstrating that the brainstem nuclei driving arousal level fluctuations are anatomically and functionally interconnected with the oculomotor nuclei that generate direction-specific activation of extraocular muscles^11–15^. Consequently, accurate interpretation requires precisely linking eye movement patterns to their underlying neurophysiology.

However, testing this hypothesis in humans has been hampered by methodological limitations in using EOG to assess oculomotor control during sleep. Specifically, the low spatial resolution and poor signal-to-noise ratio of EOG prevents the accurate measurement of saccadic kinematics like speed, which are critical for characterizing movement dynamics^16–18^. Furthermore, the EOG signal is contaminated by electrical artifacts, particularly from facial muscle activity and frontal brain activity. This presents a significant confound, as muscle artifacts can be indistinguishable from true saccades. Thus, relying solely on EOG prevents the necessary segregation of pure oculomotor events from the muscle twitches and cortical activity that accompany sleep, leaving the relationship between specific eye movement dynamics and arousal poorly defined.

To overcome these limitations, we employed in this study a multi-modal framework centered on high-resolution, infrared video-based eye-tracking. This approach allows us to establish ground truth for oculomotor behavior during sleep, providing the first comprehensive characterization of eye kinematics across all sleep stages. By bypassing the outlined limitations of EOG with computer vision, we can effectively segregate true oculomotor output from concurrent non-oculomotor sources that contaminate EOG (e.g., volume-conducted cortical activity and muscular artifacts), resolving the long-standing ambiguity between eye movements and electrical artifacts. To link these accurate oculomotor patterns to arousal, we concurrently tracked pupil size (previously reported from these same eye-tracking recordings^9^) and the aperiodic component of the EEG signal (spectral slope). This previous work has already established the feasibility of this approach and demonstrated a strong link between these arousal indices during sleep^9^. In the present study, we leverage the eye-tracking video stream to quantify eye-movement kinematics, addressing distinct questions and analyses.

Building upon our prior work, our primary hypothesis was that oculomotor activity is quantitatively coupled with—and fluctuates as a function of—pupil-based and cortical arousal across sleep macro- and microstructure. Specifically, we aimed to: (1) establish normative kinematic profiles of eye movements across sleep macrostructure and microstructural events; (2) determine the relationship between eye movement speed and concurrent indices of pupil-based and cortical arousal (pupil size and EEG spectral slope); and (3) use this approach to precisely delineate the neurophysiological differences between tonic and phasic REM sleep. Ultimately, this detailed mapping of eye positions and movements serves to validate and enrich future EOG-derived readouts, also in clinical settings.

## Methods

### Participants

Following recruitment of 18 individuals, one person was removed from the cohort due to exceeding the Ocular Surface Disease Index threshold (score > 12), indicating abnormal eye dryness^19^. The final sample consisted of 17 healthy, non-smoking adults (10 female, mean±SD age; 29.83±8.69 years) who reported a regular sleep-wake rhythm, and had a body-mass index between 17-30. Participants reported no presence of psychiatric/neurological diseases, sleep disorders, or clinically significant concomitant diseases. Ethical approval for the study was obtained from the Cantonal Ethics Committee Zurich (KEK ZH, BASEC 2022-00340), and all participants provided written informed consent before commencement, with monetary compensation provided for their involvement (CHF 70). The study was conducted in accordance with the Declaration of Helsinki.

### Experiment procedure

Prior to enrolment, a telephone screening was conducted to explain the procedure and address questions. On the day of testing, participants answered questionnaires about demographics, health status, handedness, sensitivity to noise, eye health, sleep habits, sleep quality, and daytime sleepiness (not reported here). To monitor physiological signals, we utilized gold electrodes (Genuine Grass electrodes, Natus Medical Inc., Pleasanton, US) to record EEG at positions Fpz, Cz, O1, O2, and M1 (Reference), following the international 10-20 system. Additionally, chin EMG, EOG, and ECG were tracked, ensuring all impedances remained under 20kΩ throughout the session. Subsequently, participants lay in a supine position on the bed where the sides of the pillow were wrapped with folded towels to restrict the head from side movements. Strips of Hypafix tape (BSN Medical GmbH & Co KG, Hamburg, Germany) were affixed to the upper eyelid of the right eye to maintain it open. A single strip of tape was used on the lower eyelid. To prevent eye dryness, vitamin A eye ointment was applied on the eye immediately before the eye was covered by a transparent eye bandage (PRO Optha S, Lohmann & Rauscher International GmbH & Co. KG, Rengsdorf, Germany). Finally, the eye tracker goggles (Pupil Core, Pupil Labs GmbH, Berlin, Germany) were fitted and taped firmly to the temples. Lights out (approximately 10-11pm) occurred as close as possible to the time participants reported going to bed (Fig. 1A).

**Figure 1.**
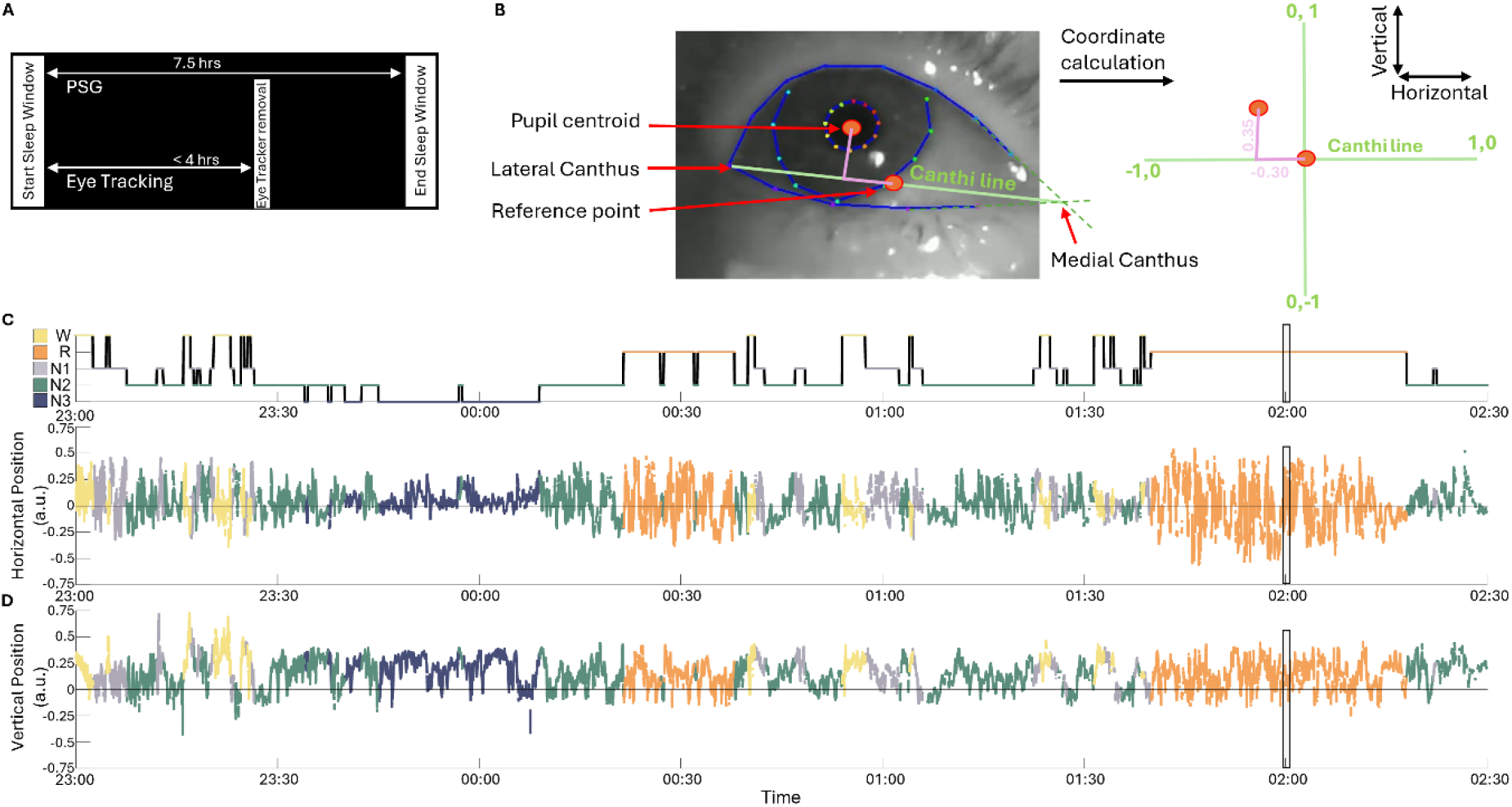
Experimental procedures. (A) Overall sleep protocol: Polysomnography was applied throughout the 7.5 hour sleep window whereas pupil size was measured only for up to the first 4 hours. (B) Calculating pupil coordinates used for eye movement analyses: Anatomical features of the eye were extracted from the videos: pupil and eyelid centroids (red markers), medial canthus (intersection of dashed green lines), canthi line (green line connecting medial and lateral canthus), reference (midpoint of canthi line). (C-J) W: Wakefulness (yellow); N1: NREM1 (gray); N2: NREM2 (green); N3: NREM3 (blue); R: REM (orange). (C-D) Hypnogram during eye tracking, along with the accompanying time series horizontal and vertical (D) pupil position coordinates, from a representative participant. The displayed time period is the first 3.5 h of a recording as shown in (A). The gray box is the time period plotted in Fig. 4D.

Except for the small infrared light source on the goggles, all light sources from the devices were covered using black tape or blackout curtains. During the complete sleep window, we recorded polysomnography and ECG (not reported here) using a BrainAmp ExG amplifier (BrainProducts GmbH, Gilching, Germany) through OpenVIBE^20^ at a sampling rate of 500 Hz. Additionally, the eye tracker video stream was recorded using Pupil Capture. The experimenter on call entered the sleep lab up to four hours from taping the eye open to remove the goggles (Fig. 1A), eye bandage, all tape strips, and the head restricting towels and participants could go back to sleep. Throughout the complete sleep window, auditory stimulation was applied using Etymotic insert earphones (Etymotic Research Inc., ER 3C). Time windows where tones were played were excluded from all analysis reported here. In the morning, the experimenter quietly entered the sleep lab, stopped all data acquisition devices, and woke the participant up. The session ended with a second round of sleep quality and mood questionnaires and reimbursement. More details on the study protocol can be found elsewhere^9^.

### EEG analysis and sleep scoring

All data prior to the moment of the horizontal and vertical eye movements (visually detected in the EOG electrodes) marking the beginning of the sleep window were excluded from further analysis. Then, EEG data was down-sampled to 200Hz using the EEGLAB^21^ toolbox function *pop_resample* in MATLAB (R2019a, MathWorks Inc., Natick, MA). Thereafter, the automatic sleep staging tool YASA^22^ was used to predict sleep stages in 30s windows. Sleep stage predictions with less than 0.6 confidence were visually inspected using the sleep stage visualizer toolbox Visbrain^23^ and manually corrected by an expert scorer following the standard AASM scoring guidelines^24^. The same procedure was applied in our previous work except here we added a step removing 30s epochs surrounding sleep stage transitions from analysis to reduce confounding effects from different neighboring sleep stages^9^.Results from figures 2 and 3 excluded 30s epochs that comprised less than 15s of valid eye-derived data. Artifacts were rejected based on a previously described semiautomatic artifact removal procedurey^25^.

**Figure 2.**
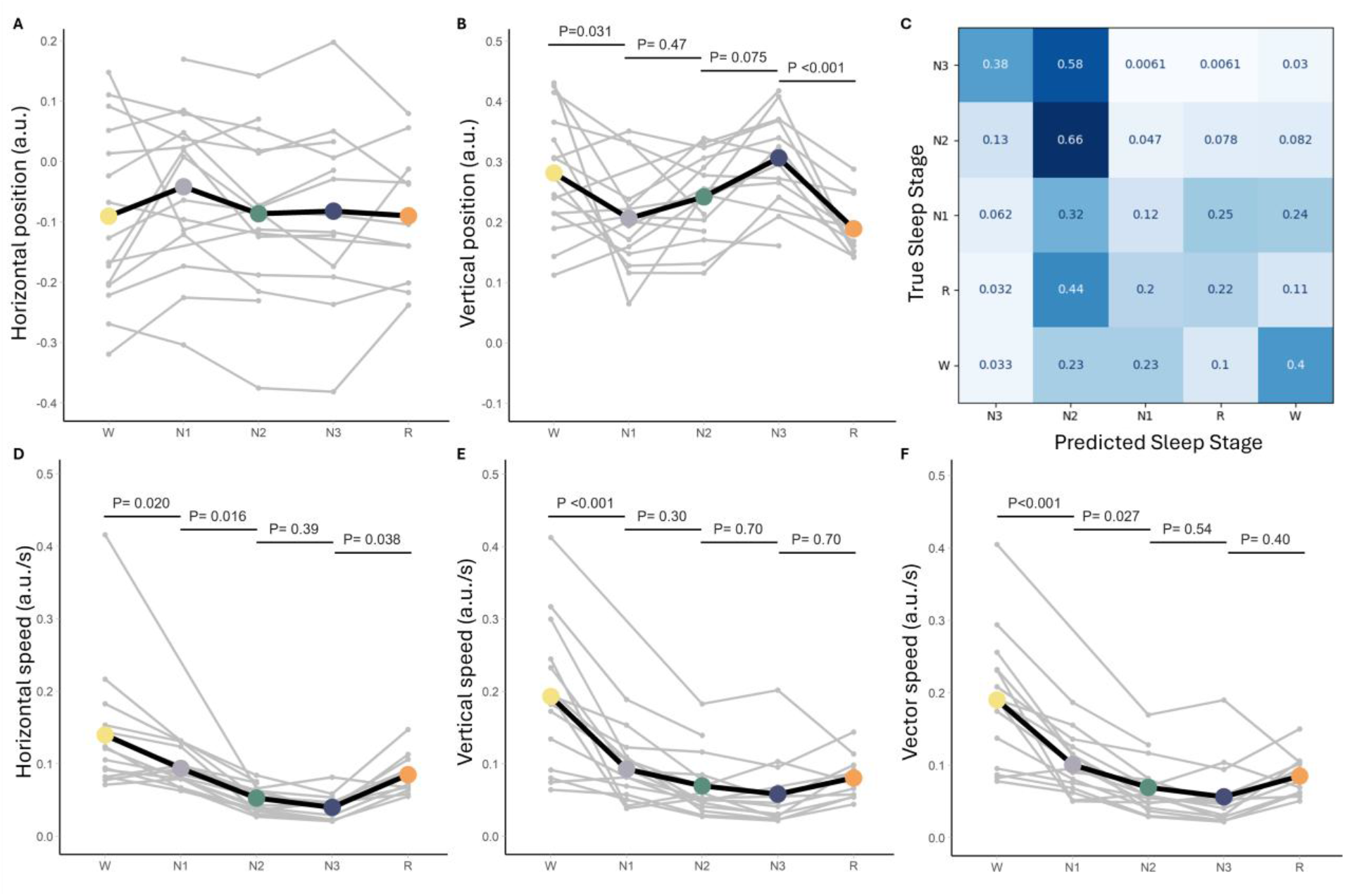
Horizontal and vertical pupil position and movement speed across sleep stages. (A) Average horizontal pupil position (F(4, 50.16)=1.59, p=0.192), and (B) vertical pupil position (F(4, 51.328)=6.90, p<0.001) for wake, N1, N2, N3, and REM sleep. (C) Confusion matrix showing the classification of sleep stages in the test set using eye-derived features (test accuracy = 47%, F1 score = 0.46). (D) Average horizontal speed (F(4, 50.681)=14.764, p<0.001), (E) vertical speed (F(4, 50.142)=25.349, p<0.001), and vector speed (F(4, 50.19)=29.837, p<0.001) for wake, N1, N2, N3, and REM sleep. The data of each participant is depicted in gray. Statistical differences between sleep stages are depicted (see supplementary tables 1-5 for more detailed information). p-values are based on post-hoc t-test and are adjusted for multiple comparisons using Benjamini-Hochberg correction for all contrasts. All statistical results are based on two-sided tests.

**Figure 3.**
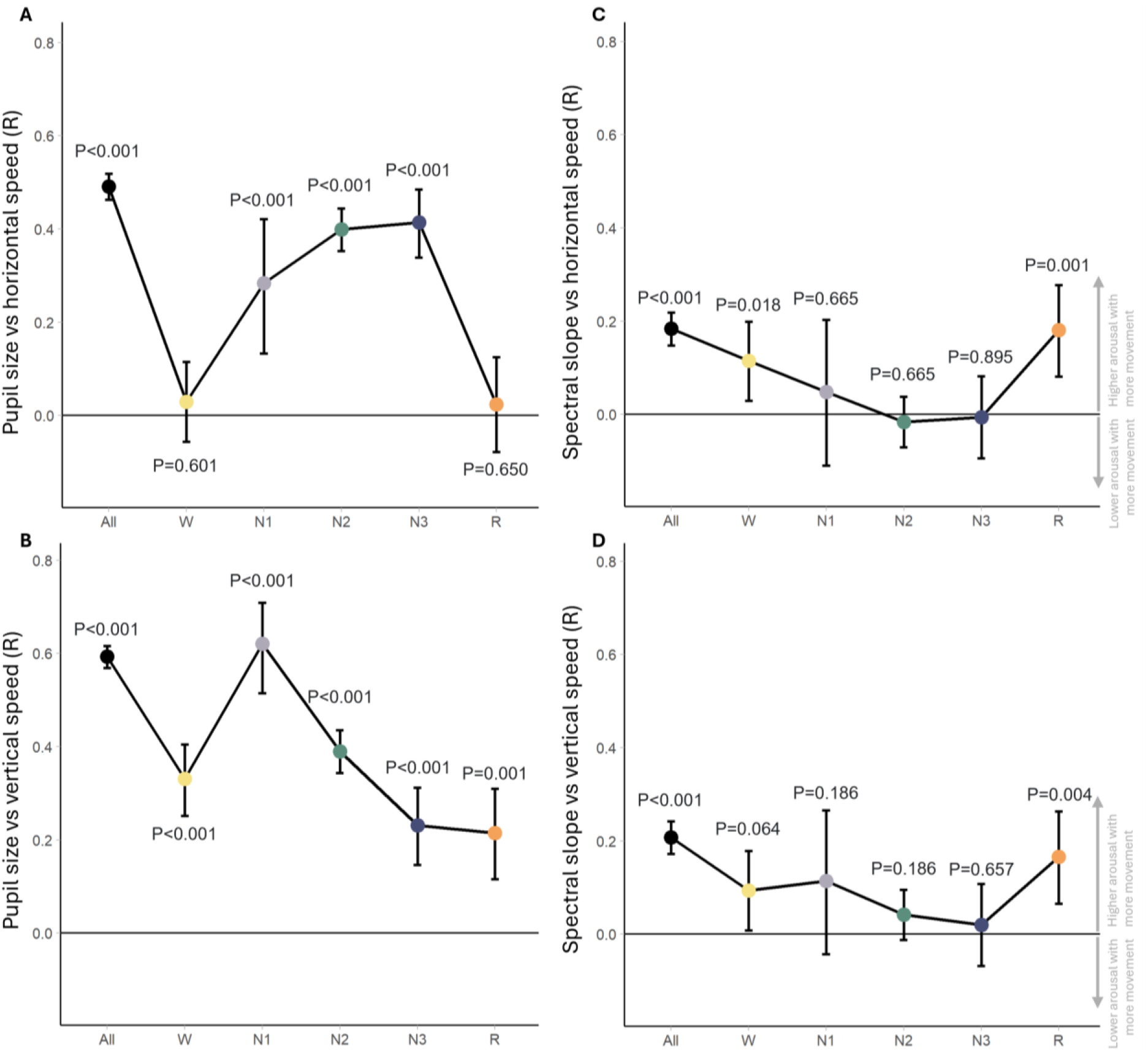
Relationship between eye movement-derived metrics and arousal metrics. (A-B) Repeated measures correlation across and within sleep stages between pupil size and horizontal speed (A) and vertical speed (B). (C-D) Repeated measures correlation across and within sleep stages between spectral slope and horizontal speed (C) and vertical speed (D). “All” refers to all sleep stages together (All: n=16, W: n=14, N1: n=15, N2: n=16, N3: n=12, R: n=11). Eye-derived measures were aggregated in 30-s segments to match the segment lengths of the spectral slope data. p-values are corrected for multiple comparisons using Bejamin-Hochberg correction. Data is presented as mean values ±95% confidence intervals.

To estimate the spectral slope, EEG signals were first 50Hz notch filtered, then 0.3Hz high-pass filtered, and finally 60Hz low-pass filtered using a Hamming windowed sinc FIR filter from the EEGLAB toolbox function *pop_eegfiltnew*. The spectral power from 0.5Hz to 45Hz was estimated using a multitaper approach^26^ which were subsequently used to extract the slope of a fitted model using the FOOOF^27^ algorithm in the 30 to 45Hz range, according to Lendner and colleagues^5^. The slope value indicates the slope of the power spectrum in the log-log scale. FOOOF settings were kept default except for minimum peak width limits set to 1 and maximum number of detectable peaks set to 4. Notably, we identified artifacts as windows with power values exceeding 3dB within the frequency range of 0.5 to 45Hz, resulting in the exclusion of 5.45% of the 30s windows derived from Fpz (Figure 3C,D).

### Offline detection of eye-derived metrics

To calculate eye-derived metrics, the eye tracker video was first resampled to 50Hz, and processed using Python 3.8 and DeepLabCut 2.2^28,29^. 36 markers were labeled in each frame: 12 marking the palpebral fissure circumference, 12 marking the iris circumference, and 12 marking the pupil circumference. Processing of the markers was conducted in MATLAB. First, the lateral and medial canthi of the right eye were derived from the palpebral fissure markers. The lateral canthus was one of the twelve markers (see Figure 1B), while the medial canthus was determined as the intersection of two lines fitted to the two most proximal markers on the upper and lower eyelids, respectively, as shown in Figure 1B. To account for palpebral fissure twitches (e.g. blink attempts during wake), the palpebral fissure markers were smoothed using a trailing average 5s moving window. The midpoint between the lateral and medial canthi served as the center coordinates of the eye position plane (horizontal, vertical = 0,0), with the line connecting the medial canthus (1,0) and the lateral canthus (-1,0) defining the horizontal axis for eye movements. Consequently, the distance between the canthi was set to 2 arbitrary units (a.u.). The vertical axis for eye movements was orthogonal to the horizontal axis, intersected at (0,0), and was scaled according to the length of the horizontal axis. Eye movement speed was calculated as the rate of change of eye position per second.

Data was then linearly interpolated using the MATLAB function *interp1* to match the 200Hz sampling rate of polysomnography data and to fill gaps with up to one second of missing data. Details on how pupil size was calculated can be found elsewhere^9^. Briefly, pupil size was determined by the average ratio of pupil to iris radii calculated from the pupil and iris markers described above. One of the 17 participants was excluded from further analysis due to insufficient valid eye position data, with only 5.64% of the 30-second epochs in which the right eye was taped open containing valid data. The remaining 16 participants had an average of 85.10% ± 23.63% of 30-second epochs with valid eye position data.

### Classification of sleep stages using eye-derived features

To determine the extent to which eye-derived metrics can independently predict sleep macrostructure, we trained a Random Forest classifier to distinguish between wake, N1, N2, N3, and REM sleep, using previously scored 30-s sleep epochs as true labels. Average horizontal and vertical position, and horizontal and vertical speed for every scored epoch were used as input features for the classifier. In supplementary analyses, pupil size was added to the feature set to evaluate its value in sleep staging. The Random Forest model (100 trees, maximum depth of 10) utilized balanced class weights, which adjusted weights inversely proportional to class frequencies in the training data to mitigate the impact of sleep-stage imbalance (e.g., there is generally an underrepresentation of N1 sleep). Thirteen participants were assigned to the training set, two to the validation set for parameter refinement, and one participant was reserved as a strictly independent test set. One participant was excluded from analysis due to low feature quality and missing data (only 5.64% of video-based eye data was valid).

Model performance was assessed via overall accuracy and weighted F1-score. To identify the specific relevance of each eye-derived metric in classifying each sleep stage, we utilized SHAP (Shapley Additive Explanations). This approach quantified the contribution of each eye-derived feature to the model’s local and global decision-making, providing a neurophysiologically interpretable ranking of how kinematics drive stage differentiation. All analyses were implemented using the *scikit-learn* and *shap* packages in Python 3.9.

### REM sleep scoring

To analyze the relationship between eye movement derived metrics and pupil-based and cortical arousal during REM sleep, we categorized REM sleep into phasic (active) and tonic (quiescent) periods. This classification utilized the norm of the horizontal and vertical coordinates of the pupil position with respect to time (i.e., vector speed in a.u./s) and was supplemented with EOG-based criteria, following previously established definitions of phasic and tonic REM sleep^10^. Phasic bouts were identified by rapid eye movements with a speed at or above 0.4 a.u./s. To capture the “burst” nature of these movements, individual events separated by at most 0.1 s were merged, and clusters occurring within 2.0 s of each other were aggregated into a single phasic bout.

Tonic REM was defined by sustained ocular quiescence where vector speed remained below 0.2 a.u./s and EOG deflections were below 25µV in both horizontal and vertical EOG channels. Only periods of such quiescence lasting at least 2.0 s were labeled as tonic. To facilitate comparison with rapidly changing arousal markers (EEG spectral slope and pupil size), identified phasic and tonic bouts were partitioned into 1.0-s non-overlapping mini-bouts. Only complete 1.0-s segments were retained for analysis, effectively linking instantaneous eye-derived metrics to the underlying neurophysiological state.

### Detection of synchronised K-complexes

We identified synchronized K-complexes to link eye-derived metrics with widespread, cortical events unrelated to the experimental stimulation protocol. K-complexes were first visually labeled in the Fpz electrode during N2 sleep, with the onset defined as 550 ms prior to the negative peak, as outlined elsewhere^9^. Importantly, only K-complexes occurring outside of the auditory stimulation windows were included, ensuring the detected events were unrelated to the experimental stimulation protocol. A K-complex was categorized as synchronized if the frontal event co-occurred with a significant deflection over central regions. Specifically, the Cz signal was baseline-corrected using the 1.0-s period immediately preceding the frontal onset. A synchronized K-complex required a negative deflection in the Cz electrode exceeding 75 µV with a peak latency within 1.0 s of the frontal onset. To provide a clean window for subsequent time-frequency analysis and ensure events were physiologically distinct, we excluded K-complexes occurring within 5.0 s of each other. This methodology allowed us to ensure that the subsequent eye-derived metric analysis was anchored to highly synchronized cortical fluctuations.

### Statistics

Statistical analyses were conducted in R (v 4.4.2; R Core Team, Vienna, Austria). Using the R packages lme4^30^ and lmertest^31^, we computed several one-factor linear mixed-effects models to investigate differences in eye-derived measures and spectral slope between sleep stages or REM state. When fixed factors were sleep stage, outcome measures included average horizontal position and speed (Fig. 2A,D), average vertical position and speed (Fig. 2B, E), average vector speed (the rate of change per second of the hypothenuse between horizontal and vertical position; Fig. 2F). When fixed factors were REM state, outcome measures included vector speed (Fig. 4A), pupil size (Fig. 4B), and spectral slope (Fig. 4C). We derived post-hoc p-values using Satterthwaite’s method from the R package lmertest and corrected for multiple comparisons with the Benjamini-Hochberg^32^. Two investigate the topographical heterogeneity of spectral slope surrounding K-complex onset, a two-factor linear mixed-effects model was computer with EEG channel and K-complex timing as fixed factors and spectral slope as outcome measure (Fig. 5E). As the two-factor linear mixed-effect model was significant for the interaction effect, we derived post-hoc p-values for the contrasts of interest using Satterthwaite’s method from the R package lmertest and corrected for multiple comparisons with the Benjamini-Hochberg method. Visual inspection of the residual plots of the linear models did not reveal any obvious deviations from normality or homoscedasticity. All mixed-effects models had participant as random factor. To investigate the relationship between eye movements and arousal within and across sleep stages, we calculated inter-participant correlations of eye movements, pupil size, and spectral slope in 30s epochs using repeated measures correlation analyses^33^. Because our dataset contains multiple observations per participant, using standard Pearson correlations would violate the assumption of independent data. P-values <0.05 were considered significant. Plots were generated using the R package^34^ and MATLAB. Where applicable, percentage values reported in the main body of text are in the format mean ± standard error of the mean.

**Figure 4.**
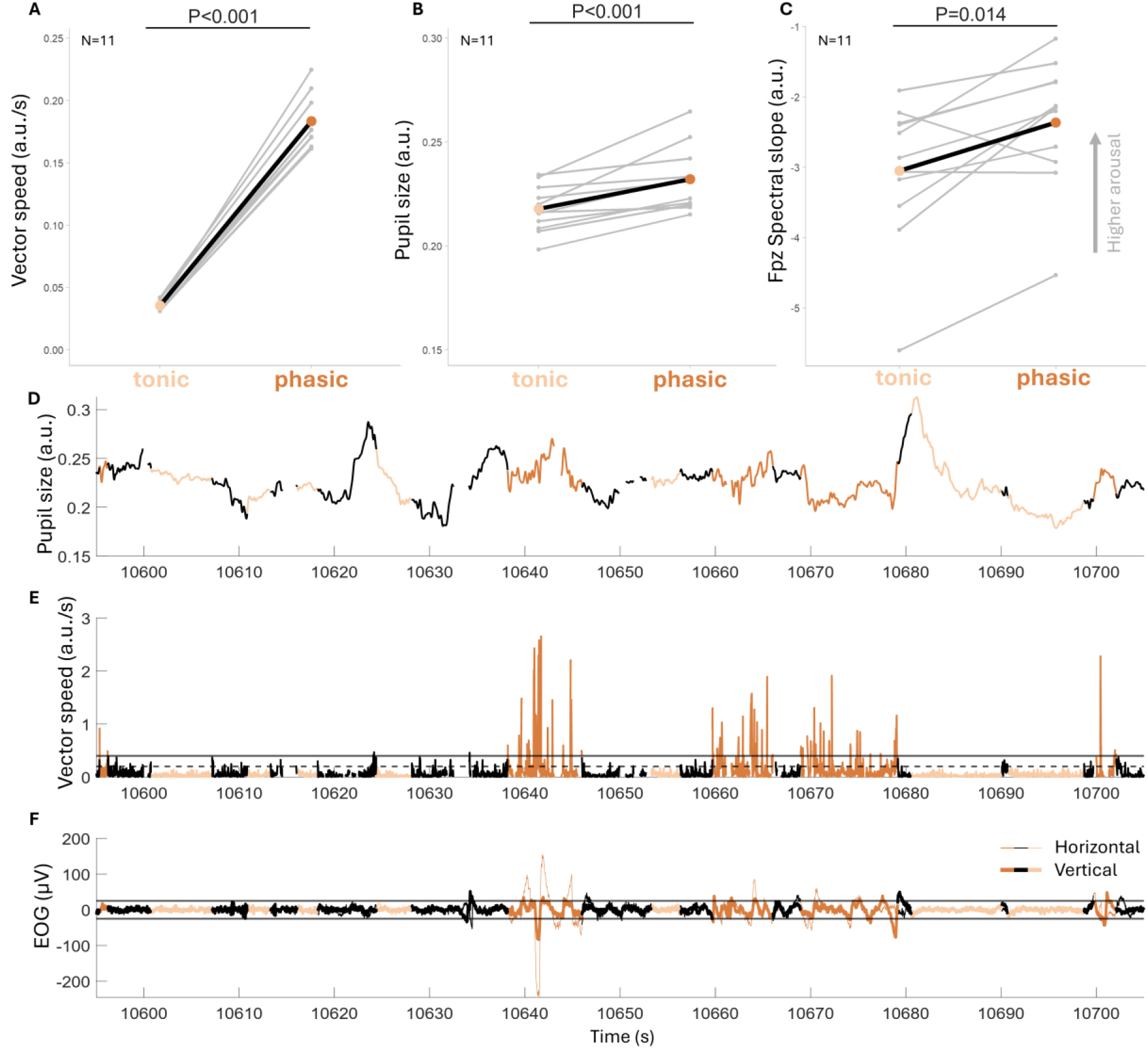
Eye movement speed, pupil size and spectral slope during tonic and phasic REM substates. (A) Vector speed, (B) pupil size, and (C) spectral slope across different EEG channels during tonic and phasic REM, respectively. (A) As expected, vector speed during phasic REM was significantly higher than during tonic REM (416.40±45.11%, p<0.001). (B) Pupil size during phasic REM was significantly larger than during tonic REM (6.56±4.40%, p<0.001). (C) Spectral slope during phasic REM was significantly flatter (less negative) than during tonic REM for the frontal electrode (Fpz: 21.25±22.97%, p=0.014). p-values are based on post-hoc t-tests and are adjusted for multiple comparisons using Benjamini-Hochberg correction. All statistical results are based on two-sided tests. (D-F): Individual example of REM pupil size (D), vector speed (E), and EOG in the horizontal derivation (thinner) line and vertical derivation (thicker line) (F), respectively. In (E), the horizontal continuous line indicates the minimum vector speed (0.4 a.u./s) for phasic REM and the horizontal dashed line below indicates the maximum vector speed (0.2 a.u./s) for tonic REM. Tonic REM also had to be accompanied by REM EOG activity within the ±25µV range depicted by the horizontal black lines. The example window in (D-F) corresponds to the time in the gray boxes in the sleep recording from Fig. 1C-D.

**Figure 5.**
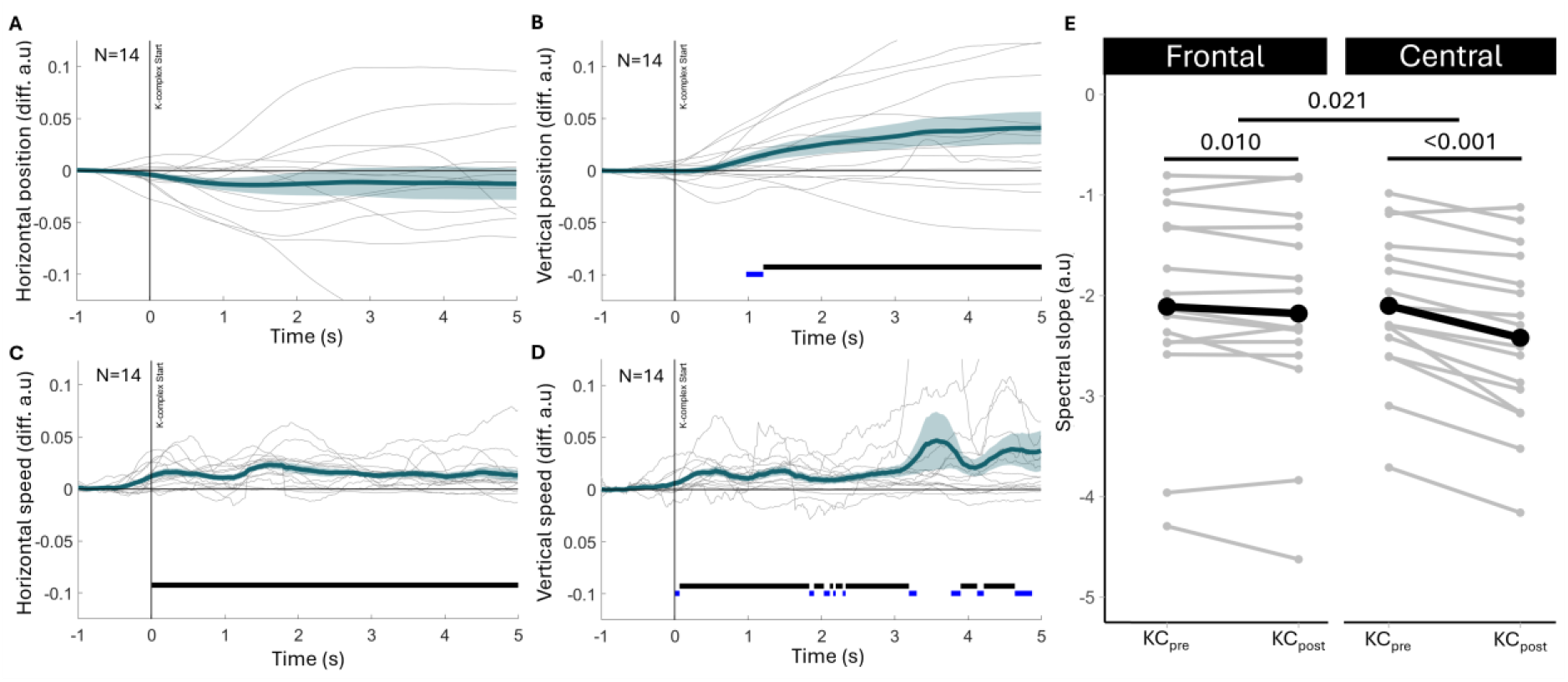
Eye movements and spectral slope during K-complexes. (A-D) Horizontal position, vertical position, horizontal speed, and vertical speed of the eye color-coded in blue and normalized to the average of -1s to -0.5s prior to K-complexes, respectively. Center lines represent the group mean and shading around the center lines the SEM. Gray lines are the mean response of each participant. Black horizontal lines mark significant differences from baseline prior to K-complexes (p<0.05). Bright blue horizontal lines mark trend-level differences from the baseline prior K-complexes (p>=0.05 and p<0.01). (E) Spectral slope in frontal (Fpz) and central (Cz) electrodes -6s to -1s before (KC_pre_) and 1s to 6s after (KC_post_) K-complexes. We found an interaction effect indicating that the increasingly steeper slopes from KC_pre_ to KC_post_ were more prominent in central electrodes than in frontal electrodes, F_kctimeXelectrode_(1, 45)=5.73, p=0.021, post-hoc comparison KC_pre_ to KC_post_ in frontal region: 5.41±8.21%. F(1, 15)=3.08, p=0.010, post-hoc comparison KC_pre_ to KC_post_ in central region: 15.38±8.01%, F(1, 15)=36.43, p<0.001. p-values are based on post-hoc t-tests and are adjusted for multiple comparisons using Benjamini-Hochberg correction. All statistical results are based on two-sided tests.

## Results

We measured pupil position and movement speed together with PSG during sleep using a novel approach we developed in healthy participants. Overall, we found that eye kinematics are highly dynamic across sleep (Fig. 1C-D).

### Pupil position and movement speed are distinct across sleep stages

EOG-derived eye movements can track large changes in pupil position but not absolute position itself. Given that EOG-derived eye movements are used to classify sleep stages, we expected to replicate with video-based eye tracking well-established eye movement speed changes between sleep stages, decreasing with deeper sleep stages until REM sleep is reached^22,24^. However, whether these eye movements are predominant in a specific axis and whether the pupil has a preferred position during sleep remains elusive. We therefore first checked whether the horizontal and vertical pupil position (Fig. 2A,B) and movement speed (Fig. 2D,E,F) calculated in 30s segments vary across sleep stages. In the horizontal axis pupil position was not significantly different across sleep stages (Fig. 2A, F(4, 50.16)=1.59, p=0.192). However, in the vertical axis the pupil was most elevated during wake and NREM stage 3 (N3) and lowest during REM sleep (Fig. 2B, F(4, 51.328)=6.90, p<0.001, see Supplementary Table 1-2). Significant differences were found across sleep stages in eye movement speed in the horizontal (Fig 2D, F(4, 50.681)=14.764, p<0.001) and vertical axis (Fig 2E, F(4, 50.142)=25.349, p<0.001), both decreasing from wake to NREM stages with horzontal eye movements becoming again faster during REM sleep when compared to NREM stage 3 sleep (see Supplementary Table 3-4). Because it remains unclear whether conventional EOG-based categorization of all sleep stages is driven primarily by vertical or horizontal movements, we calculated vector speed, defined as the rate of change in the Euclidean norm of the horizontal and vertical coordinates (Fig. 2F). Vector speed was significantly different between sleep stages (Fig. 2F, F(4, 50.19)=29.837, p<0.001), showing similar changes between sleep stages as horizontal and vertical eye movement speed (see Supplementary Table 5).

To test the distinct and rich information these four pupil position and movement speed features revealed at different sleep stages, we trained a random forest classifier with a training dataset to predict sleep stages in 30s segments on a test dataset. The classifier obtained a test performance of 47% overall accuracy and an F1 score of 0.46. The confusion matrix depicted in Figure 2C shows that the classifier had an accuracy for each sleep stage above chance level (0.2) except for NREM stage 1 (N1), the sleep stage with the the largest interscorer disagreement^35^. NREM stage 2 (N2) demonstrated the highest class-specific accuracy; however, the model showed a bias toward the majority class, frequently misclassifying other stages as N2. Based on a Shapley analysis describing the order of importance of each of the four features in labelling each sleep stage, vertical position was the most important feature to best predict REM sleep, while horizontal and vertical eye movement speed features were most important to best predict wakefulness and all NREM sleep stages (Supplementary Fig. 1D). To test whether the sleep stage classification accuracy could be improved with more video-derived features, pupil size across sleep stages was calculated as descibed in our previous research and was included as an additional input feature for the classifier (Supplementary Fig. 1A-C)^9^. The classifier’s test performance increased with this additional feature (overall test accuracy=66%, F1 score= 0.66), with a second Shapley analysis determing pupil size to be the most important feature for the classifier to best predict each sleep stage. Overall, these results show that eye-derived measures show distinct dynamics across sleep.

### Eye movements and arousal levels are distinctly correlated within different sleep stages

To assess whether the eye movement speed in distinct directions is correlated with cortical and subcortical arousal level fluctuations, we related horizontal and vertical directions of eye movement speed to pupil size and spectral slope using repeated measures correlations within each sleep stage. There were significant correlations between horizontal eye movement speed and pupil size across sleep stages (R(2900)=0.49, p<0.001, 95%CID (0.46, 0.52), with moderate correlations within NREM sleep stages (Fig. 3A, all R>=0.28, all p<0.001, all 95% CID ≥ (0.13, 0.42)). However, there was no significant relation during wake or within REM sleep (all R≤0.03, all p≥0.60). Significant correlations were also found in the vertical direction (R(2900)=0.59, p<0.001, 95%CID (0.57, 0.62) with all sleep stages showing weak to strong correlations (Fig. 3B, all R>=0.21, all p<0.001, all 95% CID ≥ (0.12, 0.30). Relating eye movement speed and spectral slope, there were significant correlations across sleep stages in the horizontal direction (R(2900)=0.18, p<0.001, 95%CID (0.15, 0.22), with weak correlations in wake and REM sleep stages (Fig. 3C, all R>=0.11, all p<0.018, all 95% CID ≥ (0.03, 0.20)). However, there was no significant relation during NREM sleep stages (all R≤0.05, all p≥0.664). Significant correlations were also found between spectral slope and eye movements in the vertical direction across all sleep stages (R(2900)=0.21, p<0.001, 95%CID (0.17, 0.24). Within sleep stages, there was only a weak correlation during REM sleep (Fig. 3D, R(370)=0.17, p=0.004, 95% CID ≥ (0.07, 0.26) all other R≤0.11, all p≥0.06).

In summary, eye movements were predominantly correlated to pupil size during NREM sleep and to spectral slope during wake and REM. This highlights that eye movements may proxy arousal levels in distinct brain regions at different moments of sleep.

### Eye-derived metrics further reveal the distinct arousal levels between REM substates

REM sleep can be categorized using EOG-derived measures into a tonic REM state with lower awakening thresholds and a phasic state with higher awakening thresholds^36^. Recent evidence showed differences in the cortical and subcortical distribution of the power spectra between these two EOG-derived substates^10,37^. However, standard EOG-based definitions of the REM substates rely on coarse temporal windows that may obscure the fine-grained, moment-to-moment arousal fluctuations driving the interplay between cortical activation and oculomotor output. To better understand arousal level dynamics between tonic and phasic REM substates, we leveraged the higher temporal sensitivity of video-based eye movements compared to EOG-based eye movements. Because it remains unclear whether conventional EOG-based categorization of tonic and phasic REM sleep is driven primarily by vertical or horizontal movements, we categorized tonic and phasic REM using vector speed (Fig. 4E-F, see Methods for details). Vector speed was significantly lower during tonic REM versus phasic REM (Fig. 4A, F(1, 10)=780.44, p<0.001). Horizontal eye movement speed did not significantly differ from vertical eye movement speed (Main Effect of Direction: F(1,40)= 0.290, p=0.5931). Furthermore, this relationship did not differ between REM substates (Interaction: F(1,40)= 1.135, p=0.2931). This highlights that our threshold-based REM substate classification criterion described in our methods section was effective in replicating EOG-based sleep stage classification without a bias to a particular eye movement direction.

Next, we investigated how cortical and pupil-based arousal levels differed between substates and whether our video-based classification of REM substates replicates previous findings^10^. Pupil size was 6.56±4.40% larger during phasic REM than during tonic REM (Fig. 4B,D, F(1, 10)=22.97, p<0.001). Spectral slope was significantly flatter (less negative) during phasic REM when compared to tonic REM independent of EEG electrode position (Fpz: 21.25±22.97%, p=0.014, Cz: 11.21±14.89%, p=0.017, O2: 11.20±11.27%, p=0.014, F(1, 10)=10.87, p<0.001, see Fig. 4C for Fpz electrode and Supplementary Fig. 3 for all three electrodes). Together, these findings highlight tonic and phasic REM substates can be categorized with video-based eye movements and that phasic REM has higher pupil-based and cortical arousal levels than tonic REM as indexed by pupil size and spectral slope, respectively.

### K-complexes are accompanied by eye movements and decreased cortical arousal

Finally, we investigated the coupling of eye movements to K-complexes, a sleep microstructural event that hallmarks changes in arousal levels. We focused our analysis on highly synchronized, pronounced K-complexes appearing in both frontal and central electrodes during NREM stage 2. By selecting these robust events, we aimed to capture microstructural markers that reflect significant fluctuations in arousal. Horizontal and vertical eye movement speed and pupil position were baseline-corrected (subtractive) to -1 to -0.5 seconds prior to K-complex onset and were plotted in a -1 to +5 second window (Figure 5A-D). We observed no systematic change in horizontal pupil position (Fig. 5A), but after onset, horizontal eye movement speed was significantly faster as compared to baseline (or 0) (Fig. 5B). In addition, after K-complex onset, the vertical position of the pupil elevated gradually (Fig. 5C) and was accompanied by a consistent increase in vertical eye movement speed when compared to baseline (Fig. 5D). These results suggest that during K-complexes the eye moves upwards while experiencing horizontal movements in a non-specific direction.

Given that we had previously found that pupil dilations, a proxy for increased subcortical arousal, accompany K-complexes^9^, we set out to investigate whether the changes in arousal levels taking place during K-complexes are also reflected in cortical arousal as indexed by spectral slope. We compared the spectral slope in frontal and central electrodes -6s to -1s before (KC_pre_) and 1s to 6s after (KC_post_) K-complexes. We found a time versus electrode interaction effect (Fig. 5E, F_kctimeXelectrode_(1, 45)=5.73, p=0.021). Post-hoc comparisons revealed steeper (more negative) spectral slopes in KC_post_ when compared to KC_pre_ most prominently in central regions (15.38±8.01%, F(1, 15)=36.43, p<0.001) and to a lesser extent in frontal regions (5.41±8.21%. F(1, 15)=3.08, p=0.010).

Together, these results show that K-complexes are accompanied by increased eye movements and lower cortical arousal as indexed by more negative spectral slope in both frontal and central cortical areas.

## Discussion

In this study, we provide a high-resolution, video-based characterization of ocular position and kinematics across human sleep, establishing a physical ground truth for eye movements that bypasses the inherent ambiguities of using electrooculography in sleep monitoring, including electrical artifacts from facial muscle twitches or frontal brain activity. By integrating eye-related metrics with measures of pupil-based and cortical arousal, we provide evidence that eye movements are not merely descriptive features of sleep stages but are quantitatively coupled to the brain’s underlying arousal state, thereby establishing a sharpened neurophysiological profile of REM substates and NREM micro-events that overcomes the limitations of electrooculography.

This ground-truth characterization reveals that the diverse dynamics of eye-derived metrics, such as higher pupil position but slower eye movement speed in NREM 3 when compared to REM and to NREM 1, are robust enough to classify sleep stages significantly above chance level, albeit with a bias towards classifying all sleep stages as NREM2 (Supplementary Table 2-3; Fig. 2). The preferred features for classification were state-dependent with horizontal eye movements being the most important feature for classifying all sleep stages except for REM, where vertical position was the most important feature (Supplementary Fig. 1A, D). Furthermore, including pupil-based arousal as a model feature substantially increased classification accuracy and restructured feature importance (Supplementary Fig. 1C). State-dependent dynamics of the eye-derived metrics and their importance in classifying sleep stages could reflect transient activity in the drivers of oculomotor and pupillary output: from rhythmic NREM fluctuations to the intense and irregular activations of REM^12–15^.

This is supported by our repeated measures correlation analyses where the coupling between eye movements and arousal metrics appears to be state-dependent. We found that within sleep stages eye movement speed correlates strongest with pupil size during NREM sleep and subsides during REM (Fig. 3A-B), where cortical arousal correlates strongest with eye speed (Fig. 3C-D) but not with pupil-based arousal (see Fig. 1I in ^9^). This repeated measures correlation analyses also confirmed the link between eye position and pupil size is biologically driven rather than a geometric artifact of the eye tracker. If movement were simply distorting the pupil measurement, we would expect consistent directional correlations across all states. Instead, we found varying correlations between speed (Fig. 3A-B) and position (Supplementary Fig. 2) such as the increase horizontal speed from NREM 3 to REM (Fig. 2D) without a maintained significant correlation with pupil size (Fig. 3A), reinforcing the validity of these metrics as independent physiological markers. Interestingly, the relationship between pupil-based arousal and eye movement speed was strongest in the vertical direction during NREM1, suggesting that Bell’s Phenomenon (or the upward rolling of our eyes as we close them) may not just be a protective mechanism but also a biomechanical manifestation of shifting brain states possibly mediated by orexin-A projections^11^.

This integrated framework of arousal proxies enables finer assessments of within-stage dynamics that traditional polysomnography often obscures, particularly regarding REM substates and K-complexes. By using video-based tracking to establish ground truth oculomotor behavior, we provide a definitive separation of tonic and phasic REM substates with high temporal resolution. Our results demonstrate that phasic REM is a distinct state characterized not only by faster eye movements but also by higher pupil-based and cortical arousal levels. Paradoxically, phasic REM has been previously described as a state of higher awakening thresholds to external sensory stimuli compared to tonic REM^36^. This discrepancy warrants further study into whether arousal levels are driven by state-dependent arousal thresholds that shift from being predominantly responsive to external stimuli, such as auditory stimuli, during tonic REM to internal stimuli, such as dream-elicited sensorimotor activity, during phasic REM^10,36,38–40^.

Another sleep microstate with a complex relationship to arousal is the K-complex, which can be viewed as either sleep protective or a reflection of transiently elevated arousal levels^1,41,42^. Our findings of a steeper post-event spectral slope (lower cortical arousal) support the former interpretation of a sleep protective cortical down-state, while the concurrent increase in eye speed and upward movement reported here, and the concurrent pupil dilation reported in our previous work, support the latter^9^. Furthermore, the accompanying eye movement dynamics raise caution in determining the extent to which the distinct drop in the EEG potential during a K-complex is driven by neuronal synchrony rather than by oculomotor changes. We report only synchronized K-complexes to further ensure that the monitored EEG signals reflect predominantly brain activity over eye movements, and the significant decrease in spectral slope after a K-complex provides further evidence of neural changes. Together, measuring proxies of various arousal level types and precisely tracking eye movements during REM and NREM microstates provide additional insights into their neurophysiological complexities and may help better identify different phenotypes of these microstates reflecting various underlying mechanisms processing external and internal stimuli.

While our 50 Hz sampling rate was sufficient for sleep staging and characterizing general kinematics, it limits the analysis of fine-grained saccadic properties. Utilizing higher sampling rates could further refine our understanding of microsaccades surrounding K-complexes and further unravel the REM state continuum proxied by eye movements. Additionally, while we have established a quantitative coupling between eye movements and arousal, the causal direction of these shifts remains unknown. The oculomotor-arousal coordination could be further elucidated by monitoring eye movements and arousal proxies while directly modulating the shared brainstem drivers^4,43–46^. This approach is particularly relevant in clinical populations such as REM Sleep Behavior Disorder, a prodromal marker of Parkison’s disease, where the coupling between the brainstem’s motor-inhibition and arousal centers fail with abnormal saccades taking place during sleep and wakefulness^47–49^. Given that video-based eye tracking offers a superior signal-to-noise ratio than EOG, complementing EOG with video-based methods in future research could help refine sleep assessments with EOG. This could then reveal novel EOG-based biomarkers of oculomotor-arousal coordination, thereby facilitating a high-fidelity re-analysis of historical EOG datasets within clinical repositories.

Our findings demonstrate that integrating oculomotor and arousal metrics provide a framework for resolving the fine-grained dynamics of sleep macrostructure and microarchitecture. By establishing this high-resolution ocular expression of the arousal continuum, we enable a more nuanced approach to sleep phenotyping that moves toward a non-discrete measurement of the sleeping brain.

## Acknowledgments

All authors thank their trainees, collaborators, and mentors for the inspiring exchanges on the thematic of this article. The authors would like to express their gratefulness to all participants of the study. This study was supported by the Swiss National Science Foundation (PZ00P3_179795 to C.L., 32003B_207719 to N.W). S.N.M. was supported by the Hochschulmedizin Zürich Flagship project STRESS. This work was also supported by ETH Zurich (22-2 ETH-026) and by the National Research Foundation, Prime Minister’s Office, Singapore under its Campus for Research Excellence and Technological Enterprise (CREATE) program (FHT).

## Author contributions

Writing – review and editing: M.C.D, S.O, T.L.O, N.W, S.N.M, C.L

Writing – original draft: M.C.D, N.W, S.N.M, C.L

Visualization: M.C.D

Data acquisition: M.C.D, S.O, T.L.O

Data analysis: M.C.D, S.O, T.L.O, N.W, S.N.M, C.L

Methodology: M.C.D, N.W, S.N.M, C.L

Conceptualization, project administration: M.C.D, N.W, S.N.M, C.L

Supervision, funding acquisition: M.C.D, N.W, S.N.M, C.L

Competing interests: The authors declare no competing interests.

