## Supplementary Material for "Video-based eye movements are linked to cortical and pupil-based arousal in human sleep"

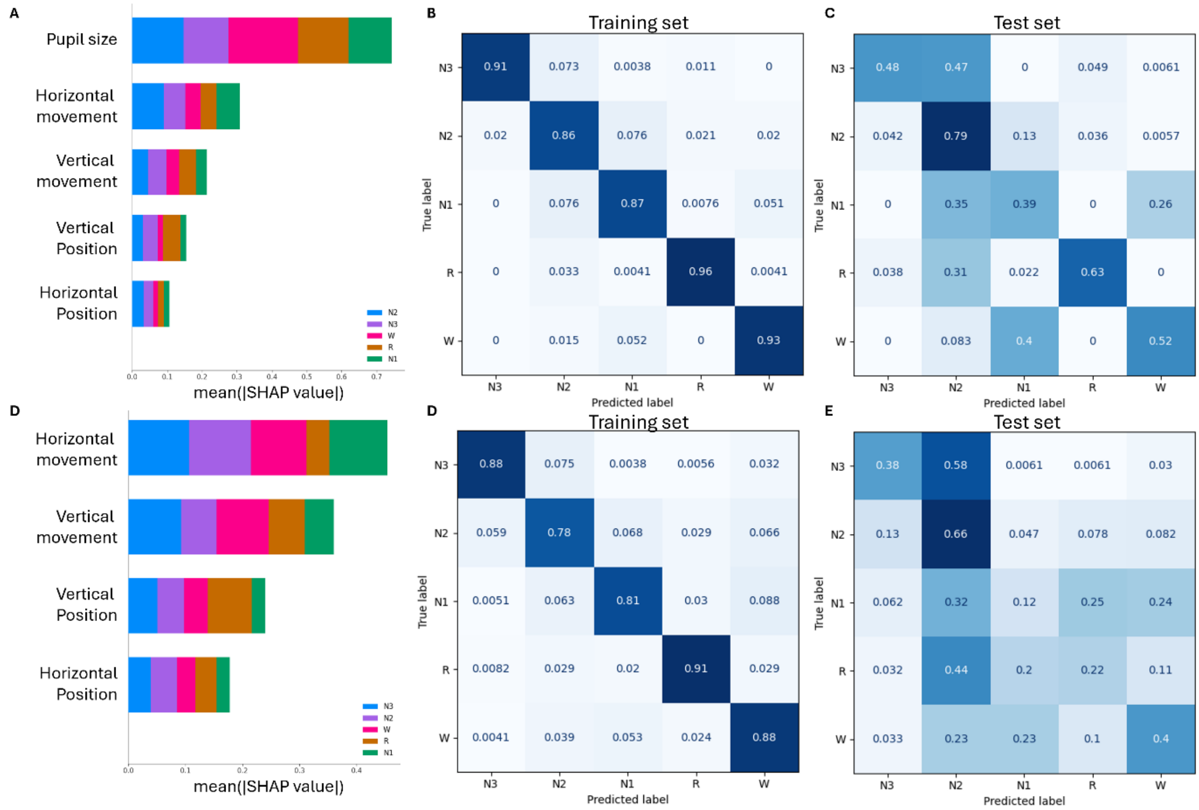


Supplementary Figure 1. Classification of sleep stages using eye-derived features.

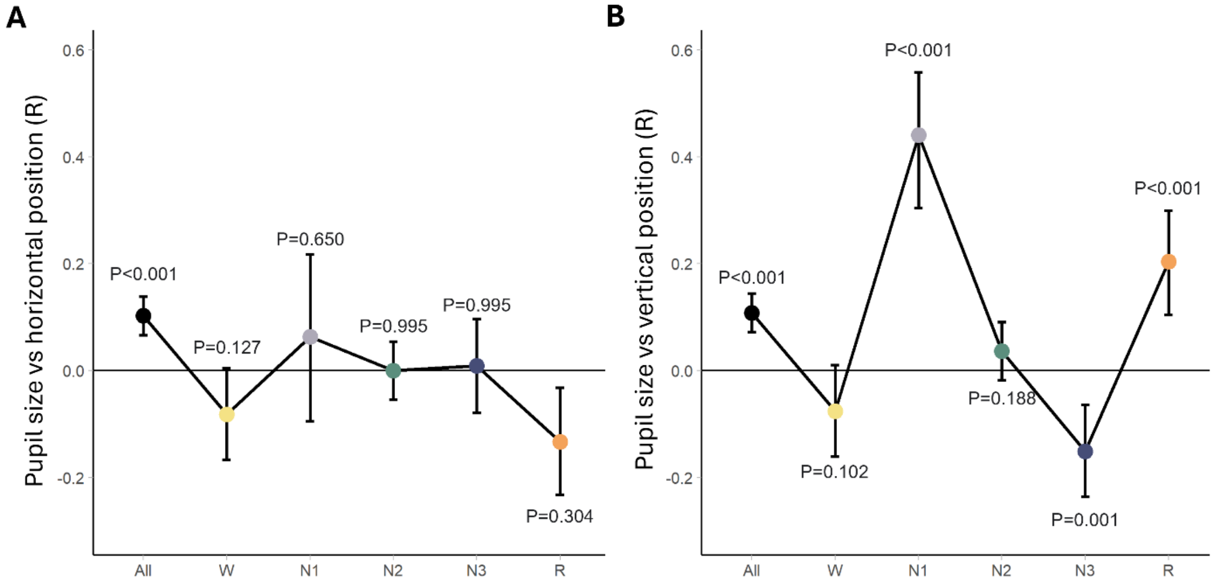


Supplementary Figure 2. Relationship between eye position and pupil size across sleep.

Repeated measures correlation within sleep stages between pupil size and horizontal position (A) and vertical position. "All" refers to all sleep stages together (All: n=16, W: n=14, N1: n=15, N2: n=16, N3: n=12, R: n=11). Pupil data was aggregated in 30-s segments to match the segment lengths of the spectral slope data. p-values are based on post-hoc t-test and are adjusted for multiple comparisons using Bonferroni correction. Data are presented as mean values ±95% confidence intervals. All statistical results are based on two-sided tests.

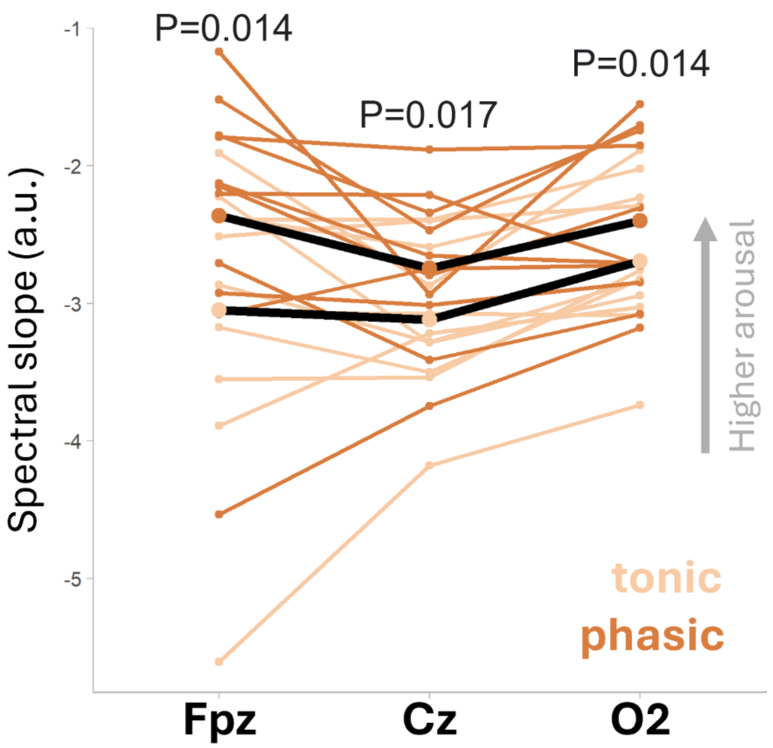


Supplementary Figure 3. Global spectral slope during REM sleep.

Spectral slope during phasic REM was significantly flatter (less negative) than during tonic REM for all three electrodes (Fpz: 21.25±22.97%, p=0.014, Cz: 11.21±14.89%, p=0.017, O2: 11.20±11.27%, p=0.014). p-values are based on post-hoc t-test and are adjusted for multiple comparisons using Benjamini-Hochberg correction. All statistical results are based on two-sided tests.

**Supplementary Table 1. Horizontal pupil position contrasts across sleep stages.** P-values are those shown in Figure 2A.

| Effect | EMMeans | Df | t.ratio | 95% CI (lower) | 95% CI (upper) | p-values | Effect size (Cohen's D) |
|---|---|---|---|---|---|---|---|
| WAKE-N1 | -0.031 | 58.396 | -1.558 | -0.09 | 0.027 | 0.873 | -0.6 |
| WAKE-N2 | 0.01 | 58.35 | 0.504 | -0.048 | 0.068 | 0.913 | 0.19 |
| WAKE-N3 | 0.014 | 58.572 | 0.675 | -0.047 | 0.076 | 0.913 | 0.27 |
| WAKE-REM | 0.008 | 58.687 | 0.336 | -0.058 | 0.073 | 0.913 | 0.14 |
| N1-N2 | 0.041 | 58.35 | 2.093 | -0.016 | 0.099 | 0.366 | 0.78 |
| N1-N3 | 0.046 | 58.572 | 2.158 | -0.016 | 0.107 | 0.351 | 0.87 |
| N1-REM | 0.039 | 58.687 | 1.737 | -0.026 | 0.104 | 0.701 | 0.74 |
| N2-N3 | 0.004 | 58.506 | 0.209 | -0.056 | 0.065 | 0.913 | 0.08 |
| N2-REM | -0.002 | 58.612 | -0.11 | -0.066 | 0.062 | 0.913 | -0.05 |
| N3-REM | -0.007 | 58.537 | -0.295 | -0.073 | 0.06 | 0.913 | -0.13 |

**Supplementary Table 2. Vertical pupil position contrasts across sleep stages.** P-values are those shown in Figure 2B.

| Effect | EMMeans | Df | t.ratio | 95% CI (lower) | 95% CI (upper) | p-values | Effect size (Cohen's D) |
|---|---|---|---|---|---|---|---|
| WAKE-N1 | 0.072 | 50.677 | 2.983 | 0.001 | 0.144 | 0.031 | 1.1 |
| WAKE-N2 | 0.039 | 50.361 | 1.635 | -0.031 | 0.109 | 0.433 | 0.59 |
| WAKE-N3 | -0.026 | 51.642 | -1.008 | -0.1 | 0.049 | 0.473 | -0.39 |
| WAKE-REM | 0.092 | 52.423 | 3.421 | 0.013 | 0.17 | 0.01 | 1.39 |
| N1-N2 | -0.033 | 50.361 | -1.406 | -0.103 | 0.036 | 0.473 | -0.51 |
| N1-N3 | -0.098 | 51.642 | -3.859 | -0.173 | -0.024 | 0.003 | -1.48 |
| N1-REM | 0.019 | 52.423 | 0.722 | -0.059 | 0.098 | 0.473 | 0.29 |
| N2-N3 | -0.065 | 51.237 | -2.589 | -0.138 | 0.009 | 0.075 | -0.98 |
| N2-REM | 0.053 | 51.955 | 2.006 | -0.024 | 0.13 | 0.251 | 0.8 |
| N3-REM | 0.117 | 51.608 | 4.271 | 0.037 | 0.198 | 0.001 | 1.78 |

**Supplementary Table 3. Horizontal pupil movement speed contrasts across sleep stages.** P-values are those shown in Figure 2D.

| Effect | EMMeans | Df | t.ratio | 95% CI (lower) | 95% CI (upper) | p-values | Effect size (Cohen's D) |
|---|---|---|---|---|---|---|---|
| WAKE-N1 | 0.043 | 50.762 | 3.017 | 0.001 | 0.085 | 0.02 | 1.11 |
| WAKE-N2 | 0.087 | 50.41 | 6.226 | 0.046 | 0.128 | <0.001 | 2.24 |
| WAKE-N3 | 0.1 | 51.81 | 6.713 | 0.056 | 0.144 | <0.001 | 2.58 |
| WAKE-REM | 0.057 | 52.681 | 3.599 | 0.011 | 0.103 | 0.005 | 1.46 |
| N1-N2 | 0.044 | 50.41 | 3.151 | 0.003 | 0.085 | 0.016 | 1.14 |
| N1-N3 | 0.057 | 51.81 | 3.829 | 0.013 | 0.101 | 0.003 | 1.47 |
| N1-REM | 0.014 | 52.681 | 0.868 | -0.033 | 0.06 | 0.389 | 0.35 |
| N2-N3 | 0.013 | 51.363 | 0.889 | -0.03 | 0.056 | 0.389 | 0.34 |
| N2-REM | -0.03 | 52.165 | -1.967 | -0.076 | 0.015 | 0.164 | -0.78 |
| N3-REM | -0.043 | 51.798 | -2.691 | -0.091 | 0.004 | 0.038 | -1.12 |

**Supplementary Table 4. Vertical pupil movement speed contrasts across sleep stages.** P-values are those shown in Figure 2E.

| Effect | EMMeans | Df | t.ratio | 95% CI (lower) | 95% CI (upper) | p-values | Effect size (Cohen's D) |
|---|---|---|---|---|---|---|---|
| WAKE-N1 | 0.093 | 50.28 | 6.348 | 0.05 | 0.137 | <0.001 | 2.34 |
| WAKE-N2 | 0.124 | 50.143 | 8.575 | 0.081 | 0.166 | <0.001 | 3.09 |
| WAKE-N3 | 0.13 | 50.76 | 8.396 | 0.084 | 0.175 | <0.001 | 3.24 |
| WAKE-REM | 0.113 | 51.099 | 6.941 | 0.065 | 0.161 | <0.001 | 2.83 |
| N1-N2 | 0.03 | 50.143 | 2.1 | -0.012 | 0.073 | 0.204 | 0.76 |
| N1-N3 | 0.036 | 50.76 | 2.344 | -0.009 | 0.081 | 0.138 | 0.9 |
| N1-REM | 0.02 | 51.099 | 1.22 | -0.028 | 0.068 | 0.699 | 0.5 |
| N2-N3 | 0.006 | 50.573 | 0.389 | -0.039 | 0.05 | 0.699 | 0.15 |
| N2-REM | -0.01 | 50.883 | -0.647 | -0.057 | 0.037 | 0.699 | -0.26 |
| N3-REM | -0.016 | 50.69 | -0.974 | -0.065 | 0.033 | 0.699 | -0.41 |

**Supplementary Table 5. Vector pupil movement speed contrasts across sleep stages.** P-values are those shown in Figure 2F.

| Effect | EMMeans | Df | t.ratio | 95% CI (lower) | 95% CI (upper) | p-values | Effect size (Cohen's D) |
|---|---|---|---|---|---|---|---|
| WAKE-N1 | 0.083 | 50.267 | 6.249 | 0.044 | 0.122 | <0.001 | 2.3 |
| WAKE-N2 | 0.121 | 50.136 | 9.283 | 0.082 | 0.159 | <0.001 | 3.35 |
| WAKE-N3 | 0.129 | 50.726 | 9.281 | 0.088 | 0.17 | <0.001 | 3.58 |
| WAKE-REM | 0.106 | 51.05 | 7.219 | 0.063 | 0.149 | <0.001 | 2.95 |
| N1-N2 | 0.038 | 50.136 | 2.909 | 0 | 0.076 | 0.027 | 1.05 |
| N1-N3 | 0.046 | 50.726 | 3.325 | 0.005 | 0.087 | 0.01 | 1.28 |
| N1-REM | 0.023 | 51.05 | 1.589 | -0.02 | 0.066 | 0.404 | 0.65 |
| N2-N3 | 0.008 | 50.547 | 0.618 | -0.032 | 0.048 | 0.539 | 0.23 |
| N2-REM | -0.014 | 50.843 | -1 | -0.057 | 0.028 | 0.539 | -0.4 |
| N3-REM | -0.023 | 50.658 | -1.52 | -0.067 | 0.021 | 0.404 | -0.63 |
